# Multi-particle cryo-EM refinement with *M* visualizes ribosome-antibiotic complex at 3.7 Å inside cells

**DOI:** 10.1101/2020.06.05.136341

**Authors:** Dimitry Tegunov, Liang Xue, Christian Dienemann, Patrick Cramer, Julia Mahamid

## Abstract

Cryo-electron microscopy (cryo-EM) enables macromolecular structure determination *in vitro* and *in situ*. In addition to aligning individual particles, accurate registration of sample motion and 3D deformation during exposures is crucial for achieving high resolution. Here we describe *M*, a software tool that establishes a reference-based, multi-particle refinement framework for cryo-EM data and improves the results of structure determination. *M* provides a unified optimization framework for both *in vitro* frame series and *in situ* tomographic tilt series data. We show that tilt series data can provide the same resolution as frame series, indicating that the alignment step no longer limits the resolution obtainable from tomographic data. In combination with Warp and RELION, *M* improves upon previous methods, and resolves a 70S ribosome bound to an antibiotic inside bacterial cells at a nominal resolution of 3.7 Å. Thus, computational tools are now available to resolve structures from tomographic *in situ* cryo-EM data at residue level.

## INTRODUCTION

Single-particle analysis^1^ (SPA) is the image analysis technique that established cryo-electron microscopy^2^ (cryo-EM) as a widely used method for macromolecular structure determination^3, 4^. In the SPA workflow, many noisy 2D observations of macromolecular particles made in a transmission electron microscope (TEM) are iteratively aligned, classified and averaged to reconstruct high-resolution 3D maps of the macromolecules’ Coulomb potential^1^. A central assumption in SPA is that each image shows a single particle in isolation, and can thus be analyzed independently of other particles.

Typical cryo-EM micrographs capture a multitude of macromolecular particles embedded in a layer of amorphous ice. As the sample is irradiated with electrons, mechanical instrument instabilities and beam-induced motion (BIM) lead to changes in particle positions and orientations throughout the exposure^5^. If left uncorrected, these changes decrease the apparent image quality and limit the map resolution. Exposure fractionation into multiple frames captures the particles at several steps along their trajectories. At fine enough fractionation and accurate motion registration, the detrimental effects of sample movement can be reversed to obtain better maps^6, 7^. Unfortunately, the granularity of the motion model is limited by the low signal per particle. Although each particle’s trajectory is unique, correlations between particles exist on a local scale and can be used to regularize the motion model^6, 8^. It is thus beneficial to treat the contents of a micrograph as a physically connected multi-particle system rather than isolated particles.

Two types of cryo-EM data are typically analyzed to obtain high-resolution maps: First, *in vitro* samples of proteins are prepared at concentrations where individual particles can be distinguished in 2D projections, and fractionated exposures at constant stage orientation (“frame series”) are acquired. Second, *in situ* samples that capture portions of crowded cellular environments, or samples containing multiple particles in close proximity and/or stacked along the projection axis, require a tomographic approach to distinguish the particles in 3D. To achieve this, the microscope stage is tilted to different angles between sub-exposures (“tilt series”). Within one tilt series, each sub-exposure usually also comprises a frame series (called here a “tilt movie”). Mechanical instability requires the stage position and orientation at each angle to be introduced as additional variables during processing. Thus, frame series can be treated as a special case of tilt series, and approaches designed for tilt series can be used for frame series as well. Map refinement from tilt series data is often called “sub-tomogram averaging” due to its use of intermediate 3D reconstructions for each particle^9, 10^ instead of 2D images. Because the refinement of both data types falls conceptually under the SPA umbrella, and because the approach we introduce does not use sub-tomograms, we use “SPA” for both cases here.

At the data pre-processing stage, when no particle positions and reference maps are available, the sample motion model can be fitted based on raw data alone using reference-free approaches^6, 7, 11-13^. Frame series are aligned and averaged in 2D, whereas tilt series are aligned and used to reconstruct tomograms. The results are fed into an SPA pipeline to obtain 3D references. A reference-based algorithm can then be used to improve the motion model accuracy by aligning individual particle frames or tilts to high-resolution reference projections. Such algorithms exist for both frame and tilt series data^8, 14, 15^, and most of them improve the accuracy by enforcing local smoothness between particle trajectories on different spatio-temporal scales. Although most motion registration algorithms exploit the multi-particle nature of the data in some way, their implementations remain different for frame and tilt series data, and limited to only one reference species even in highly heterogeneous data sets. Furthermore, they are decoupled from other parts of the refinement process, such as rotational alignment and contrast transfer function (CTF) fitting. This leads to a fragmented workflow and decreased convergence speed, potentially limiting the final map resolution.

Here we present *M*, a software tool that integrates reference-based refinement of particle motion trajectories with other parts of the SPA pipeline in a user-friendly manner. We formulate our approach explicitly in a multi-particle framework, which allows us to unify the processing of frame and tilt series, define a set of intuitive regularization constraints, and include any number of particle species at different resolutions. Coupled with a robust approach to CTF correction and with neural network-based map denoising, *M* achieves higher resolution on several exemplary frame series and tilt series data sets compared to other methods^14-17^. We demonstrate how various features of *M* contribute to these improvements, and achieve the same high resolution for frame series and tilt series data given similar amounts of particles. Most strikingly, we use *M* to visualize a 70S ribosome bound to an antibiotic in its native cellular context at residue-level resolution from *in situ* tilt series data.

## RESULTS

### Overall design

*M* was designed to form the last part of a largely automated cryo-EM data pre-processing and map refinement pipeline – preceded by Warp^12^ and RELION^18^, or compatible tools (Fig. 1). Warp performs initial, reference-free motion correction and CTF estimation on frame series or tilt movies during data acquisition. For tilt series pre-processing, Warp, starting with version 1.1.0, automatically calls routines from IMOD^19^ to perform the initial tilt series alignment, estimates per-tilt CTF using the tilt angles as constraints, and reconstructs the tomographic volumes at a large pixel size for visual analysis and particle picking. Warp then picks the particles using a convolutional neural net-based (CNN) approach for frame series, or template matching for frame or tilt series, and exports them as images or reconstructed volumes depending on the data type. In case of tilt series, 3D CTF volumes containing the missing wedge and tilt-dependent weighting information are generated for each particle^20^. The particles are then refined and classified in RELION using a multitude of strategies available there^21^. All classes and their respective refinement results are finally imported into *M* to perform a more accurate, reference-based, multi-particle frame or tilt series refinement and obtain the final high-resolution maps. Optionally, the refined parameters can be used to re-export more accurately aligned particles for further classification in RELION or compatible software. The new alignments can be applied to generate 3D volumes at higher resolutions to be used for further particle picking.

**Figure 1.**
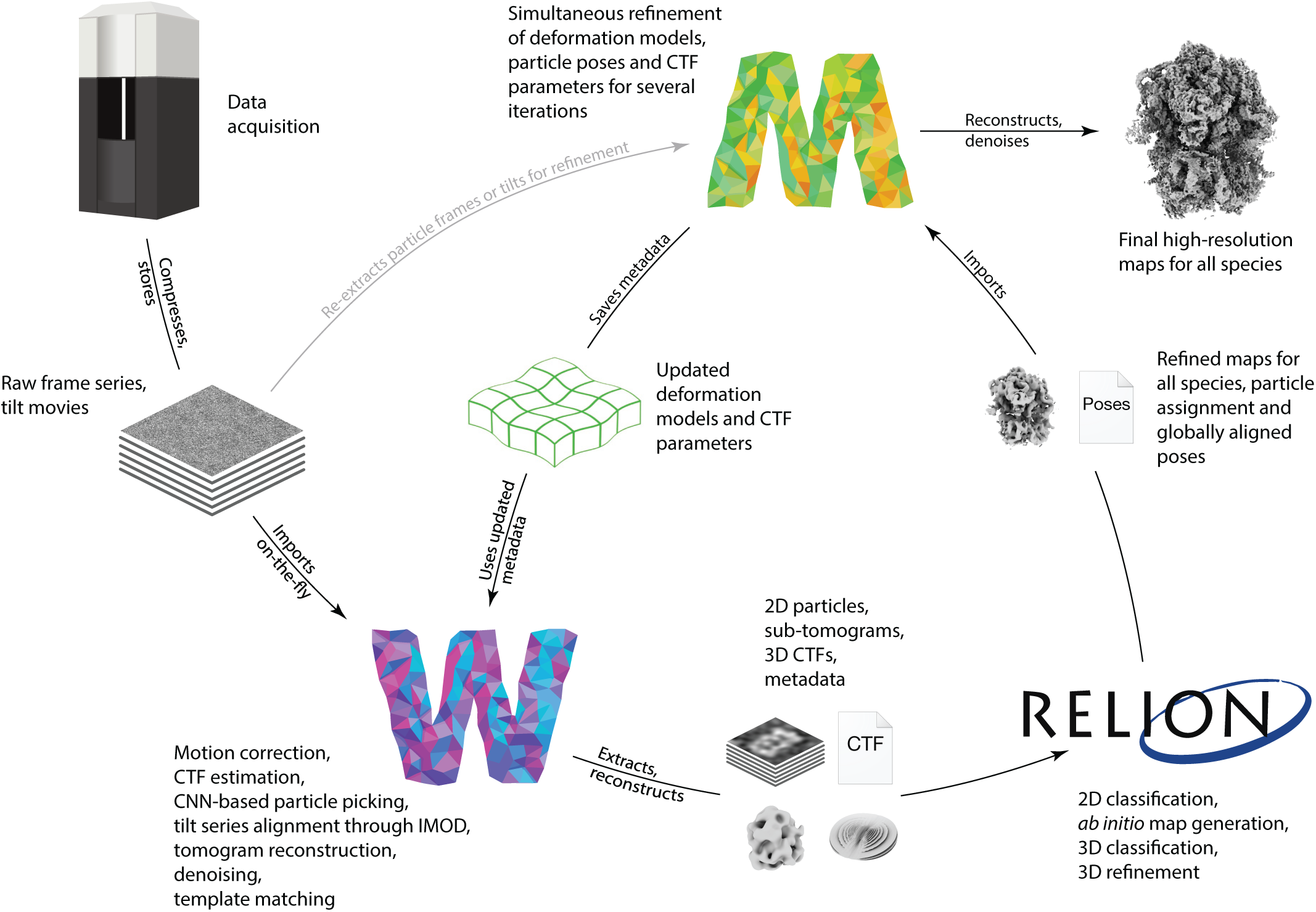
The Warp–RELION–*M* pipeline for frame and tilt series cryo-EM data refinement. Electron microscopy data are pre-processed on-the-fly in Warp, which then exports particles as images or sub-tomograms. Particles are imported in RELION, where they can be subjected to a multitude of processing strategies, resulting in 3D reference maps, global particle pose alignments, and class assignments. The particle population encompassing all classes is then imported in *M*, where reference-based frame or tilt image alignments are performed simultaneously with further refinement of particle poses and CTF parameters to improve resolution. Finally, *M* produces high-resolution reconstructions that can be used for model building. Alternatively, the improved alignments can be used in Warp to re-export particles for further, more accurate classification in RELION.

*M* provides a graphical user interface (GUI) that allows the user to create, import, export and manage data. Projects are organized as “populations”, which contain “data sources” and “species”. A data source is a set of frame or tilt series, that stem ideally from the same sample grid and acquisition session. A species is any distinct type of macromolecule, or its compositional and conformational sub-state. Each version of a data source or species is tracked using a cryptographic hash of its current state, the preceding version, and the processing parameters connecting them. The entire refinement evolution can be tracked as a directed graph, parts of which can be stored in different locations while remaining uniquely connected through the hashes. Thus, multiple users can process different collections of data sources and species independently and merge them later. This is especially useful for processing *in situ* data sets, where each user may process only a small fraction of all species contained and everyone would benefit from pooling data and results together.

### Multi-particle system modeling

*M* considers the entire field of view of a frame or tilt series as a physically connected system of particles (Fig. 2a). The particles can belong to different refined species, which can be of varying size, symmetry, and resolution. As parts of the same system, the particles are subject to the same global transformations such as the translation and rotation of the microscope stage, as well as locally similar transformations caused by BIM that result in apparent translation and rotation of particles. *M* performs a reference-based registration of these transformations (Fig. 2b), and reverses them when back-projecting individual particle frame or tilt images to obtain more accurate reconstructions.

**Figure 2.**
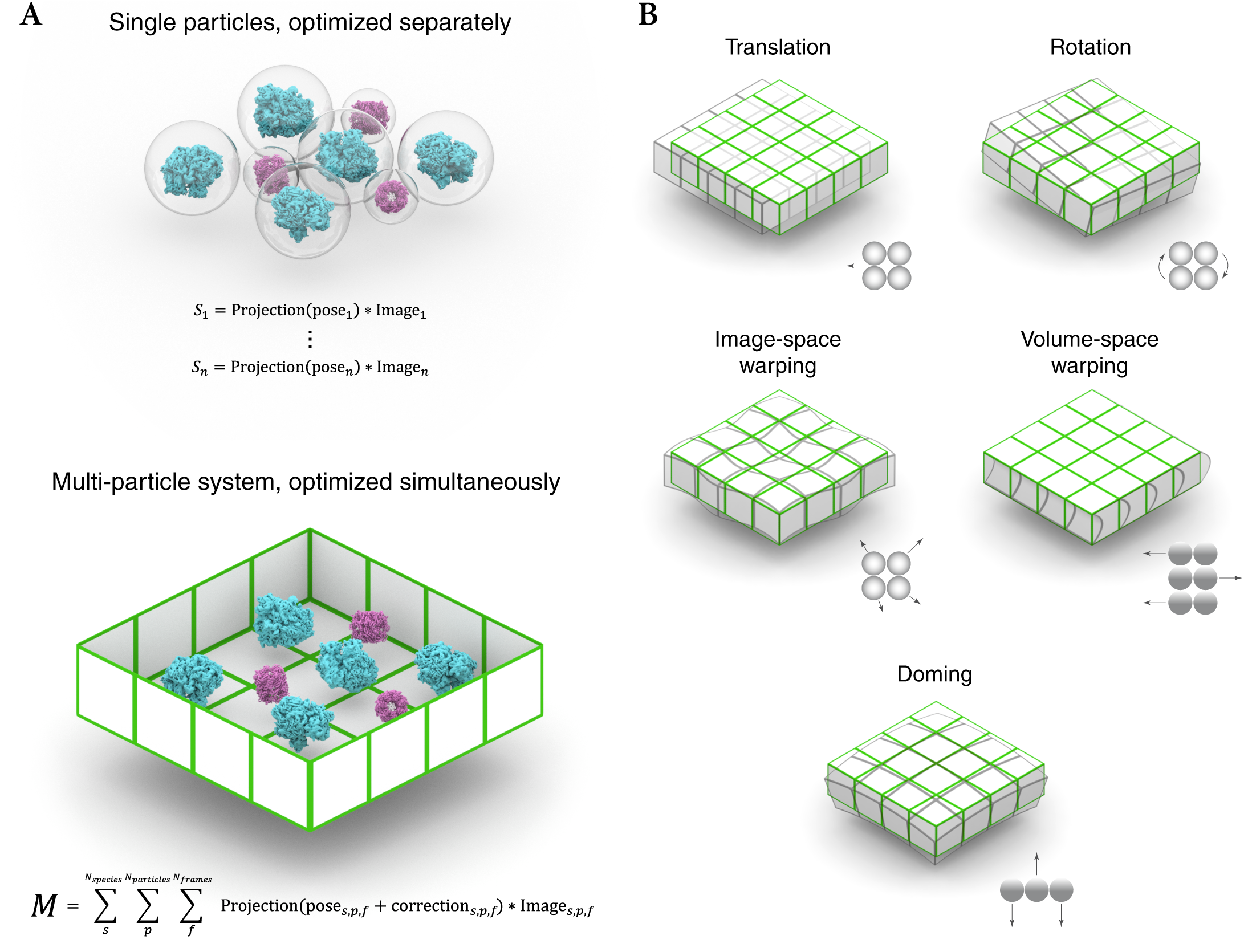
Multi-particle system modeling and optimization *M* employs a reference-based multi-particle optimization to model sample deformation and improve map resolution. (a) Particles are treated as isolated entities in SPA. Each particle has its own cost function based on the similarity between a simulated reference projection and the experimental particle image, which is optimized independently. However, particles in a real sample are physically connected and experience locally similar effects during exposure. Each imaged location is modeled as a multi-particle system. Its state model is fitted using a single cost function, which compares simulated reference projections to all experimental particle frame or tilt images. The particle poses in each frame or tilt are additionally affected by the modeled deformation of the multi-particle system, which is optimized together with the per-particle pose alignments. (b) The multi-particle system deformation model incorporates several modes: Global movement and rotation to account for inaccuracies in stage movement between frames and stage rotation between tilts; image-space warping to model local non-linear deformation in the 2D reference frame of a frame or tilt image; volume-space warping to model the movement of overlapping particles perpendicular to the projection axis (tilt series only); doming to account for the hypothesized bending of a thin sample along the projection axis (frame series only).

In frame series, all transformations occur in the same image reference frame. Their combined effects are parametrized as a pyramid of 3D cubic spline grids, where the top grid has low spatial and per-frame temporal resolution, and subsequent grids have increased spatial and decreased temporal resolution. This model is similar to the one used in Warp but fits more parameters due to the higher accuracy of reference-based registration. The user can set the spatial resolution of the top grid to adjust the model’s resolution to the available signal. In addition to image-space warping, *M* can fit doming-like motion that is known to occur at the beginning of an exposure^5^. This is implemented as parameter grids for defocus and orientation offsets with 3×3 spatial and per-frame temporal resolution.

In tilt series, *M* distinguishes image-space and volume-space effects because the tilt images show the volume from different angles. Image-space transformations are parametrized as a 3D cubic spline grid with per-tilt temporal, and a spatial resolution set by the user depending on the available signal. Additionally, parameters of a coarse 3D cubic spline grid can be fitted for every tilt movie to account for the significant exposure and deformation captured in each of them. Volume-space transformations, such as the shearing of a thick sample, are modeled as a 4D parameter grid with quadrilinear interpolation, with the accumulated exposure as the temporal dimension. Because *M* does not average particle frames or tilts in intermediate steps, per-particle translation and rotation trajectories can be fitted to model the most local transformations. The temporal resolution of the trajectories can be set for each species depending on its size and thus the signal available per particle.

To test the various features of *M*, we collected cryo-EM frame and tilt series data on an apoferritin sample (data sets AF-f and AF-t, see Methods). Using the frame series data, we show the benefit of considering the particles of multiple species in refinement. To this end, we artificially split the apoferritin population in 2 species comprising 5% or 95% of the particles (Fig. S1a). No structural similarity between the two species was assumed during refinement. Refining the 5% species alone produced a 3.2 Å map, while adding the 95% species to the multi-particle system improved the map calculated from the 5% species to 2.8 Å (Fig. S1b). This demonstrates that our multi-species refinement approach can improve the resolution for heterogeneous data sets.

### Correction of electron-optical aberrations

In addition to a geometric deformation model, *M* fits CTF parameters and higher-order aberrations including beam tilt. For frame series, defocus is optimized per-particle, similar to cisTEM^16^ and recent RELION versions^17^. For tilt series, defocus is optimized per-tilt, similar to the capability offered in emClarity^14^. For both types of data, astigmatism, anisotropic pixel size and higher-order aberrations are fitted and corrected per-series.

CTF correction at high defocus can introduce artifacts if the chosen particle box size is too small to retain high-resolution Thon rings, leading to their aliasing (Fig. S2a) and limiting the resolution for many combinations of high defocus images and small particles. *M* automates the selection of a sufficiently large box size at which the data are pre-multiplied by an aliasing-free CTF. The images are then cropped in real space. To match the underlying CTF of these images, correctly band-limited CTF^2^ images are constructed in a similar way (Fig. S2a). Both are then used for refinement and reconstruction.

We show the benefit of this approach by reconstructing a map from a previously refined high-defocus tilt series of HIV1 virus-like particles (“TS_01”, EMPIAR-10164). Using twice the particle diameter as the box size, the resolution is limited to 3.9 Å as the average sign error of the aliased CTF increases (Fig. S2b). Pre-multiplying the data and CTF at an aliasing-free size and then cropping them significantly improved the resolution to 3.2 Å using the same reconstruction box size. Only pre-multiplying the data but using an aliased analytical CTF^2^ for the Wiener-like reconstruction filter did not decrease the nominal resolution in this case. However, for algorithms that would use such aliased models during refinement and classification as well, we expect these effects to be more noticeable. This approach improved the empirically estimated per-tilt series weighting factors (see Methods) for high-defocus data to the level of low-defocus data for the entire EMPIAR-10164 data set (Fig. S2c).

### Optimization procedure

*M* optimizes all selected hyperparameters describing geometric deformation (Fig. 2b), electron-optical aberrations, and particle pose trajectories, simultaneously. Because exhaustive search over an ensemble of thousands of parameters would be impossible, *M* performs a local, gradient descent-type optimization using the Limited-memory Broyden–Fletcher–Goldfarb–Shanno (L-BFGS) algorithm^22^. To compute the derivatives for the variables efficiently, *M* precomputes sets of weights using a strategy similar to Warp’s^12^ (see Methods). Derivatives for many of the parameters can then be computed as weighted sums of per-particle image derivatives, which in turn are calculated using GPU-accelerated routines. The optimization procedure considers the signal of all defined particle species simultaneously to maximize the particle density in each frame or tilt series, thus increasing the hyperparameter fitting accuracy for heterogeneous data sets.

At the end of an optimization iteration, similar to the Fourier Ring Correlation (FRC) approach introduced previously^23, 24^, *M* calculates the per-Fourier component normalized cross-correlation between reference projections and image data. This can be used to empirically optimize anisotropic exposure-and tilt-dependent data weighting, and reconstruct new half-maps using the updated model, correcting for Ewald sphere curvature^25^. The denoiser routine is based on deep learning and is re-trained on the new half-maps (Fig. S3; see Methods). Then, various map metrics, including global, local, and anisotropic resolution, are calculated. Further optimization iterations can be performed to arrive at a denoised or low-pass filtered and sharpened map. Alternatively, 2D particles or sub-tomograms can be extracted and reconstructed from the raw data using the updated alignment information, to be exported to RELION for further, more accurate classification.

### Contribution of different model parameters to map resolution

We used the apoferritin frame and tilt-series data sets, collected from the same grid square under identical conditions (see Methods), to estimate the contribution of different groups of optimizable parameters to the quality of the reconstructed maps (Fig. 3). For frame series, particles extracted following reference-free alignment in Warp and refined in RELION (without polishing and CTF refinement) provided a baseline resolution of 2.75 Å, which was then improved by accumulating the following sets of optimizable parameters in *M*: Reference-based global motion alignment improved the resolution to 2.73 Å. Relaxing this constraint to allow local motion alignment improved the resolution to 2.71 Å. Resolving individual particle pose trajectories as a function of exposure led to a resolution of 2.66 Å. Fitting per-particle defocus and per-frame series astigmatism and beam tilt improved the resolution to 2.45 Å. Data-driven anisotropic weight estimation improved the resolution to 2.39 Å. Finally, resolving doming-like motion slightly improved the resolution to 2.32 Å.

**Figure 3.**
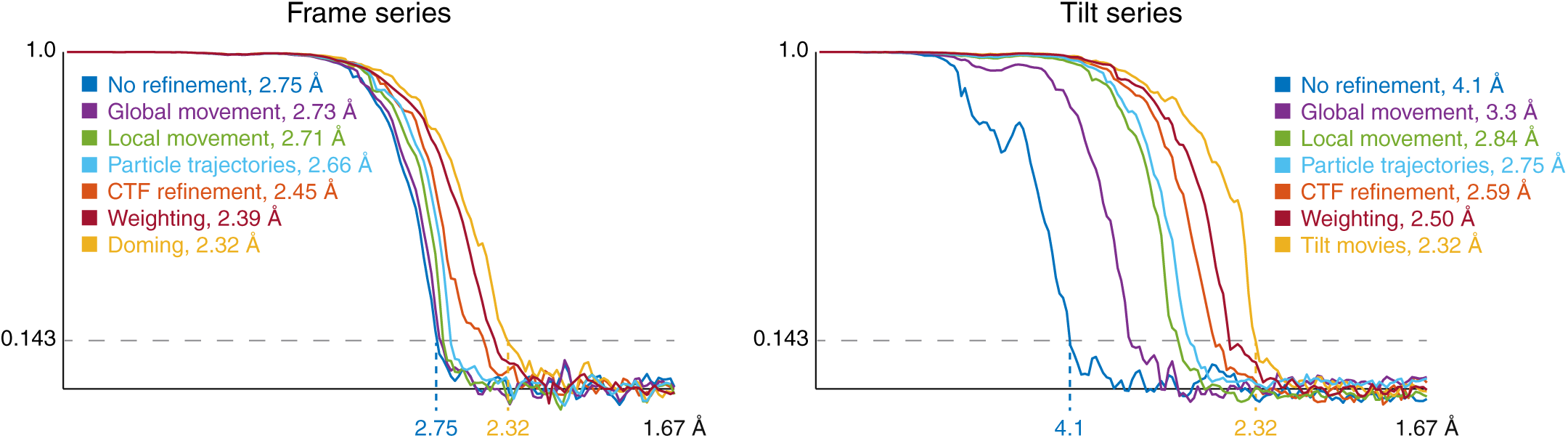
Contributions of individual multi-particle system model components to map resolution. Fourier shell correlation between half-maps for frame series and tilt series apoferritin data obtained through extending the set of optimizable parameter groups. Starting with the ‘No refinement’ baseline, in top-down order in the legend, a new group of parameters was added, while keeping the previously added groups, and refinement was performed from scratch. The resolution for each step is given in the legend.

For tilt series, sub-tomograms reconstructed following reference-free tilt movie alignment in Warp, patch tracking-based tilt series alignment in IMOD and refinement in RELION provided a baseline resolution of 4.1 Å, which was then improved by accumulating the following optimizations in *M*. First, reference-based global tilt image alignment improved the resolution to 3.3 Å. Relaxing this constraint to allow local image-space warping improved the resolution to 2.84 Å. Resolving individual particle poses as a function of exposure increased the resolution to 2.75 Å. Fitting per-tilt defocus and astigmatism, and per-tilt series beam tilt improved the resolution to 2.59 Å. Data-driven anisotropic weight estimation improved the resolution to 2.50 Å. Finally, reference-based tilt movie alignment led to a resolution of 2.32 Å. Volume-space warping was not tested because the particles were arranged in a single 2D layer.

From these tests, we conclude that accurately registering image-space deformation is essential for obtaining high-resolution maps from frame and tilt series data, whereas modeling other effects leads to smaller improvement that may generally only become significant in the sub-5 Å resolution range. Initial reference-free alignment is significantly less accurate for tilt series than for frame series. However, it is accurate enough to obtain initial reference maps and particle poses that can be further refined in *M*.

### Comparison between frame and tilt series performance

Because tilt series are often associated with lower resolution compared to frame series, we processed both types of data collected from grid holes in close proximity during the same data acquisition session on an apoferritin sample (data sets AF-f and AF-t, see Methods) to test potential intrinsic limitations of the tilt series data. Given equal amounts of particles, *M* was able to achieve the same resolution with very similar map features (Fig. 4) for both frame series and tilt series data. Thus, collecting data as tilt series does not incur a resolution penalty. However, because tilt series data are still much slower to acquire^26^ and commonly used for more crowded, thicker samples, we expect maps derived from tilt series data to remain at lower resolution on average. Nevertheless, our results show that there is no intrinsic resolution loss when using tilt series over frame series when enough high-quality data are collected and optimally processed.

**Figure 4.**
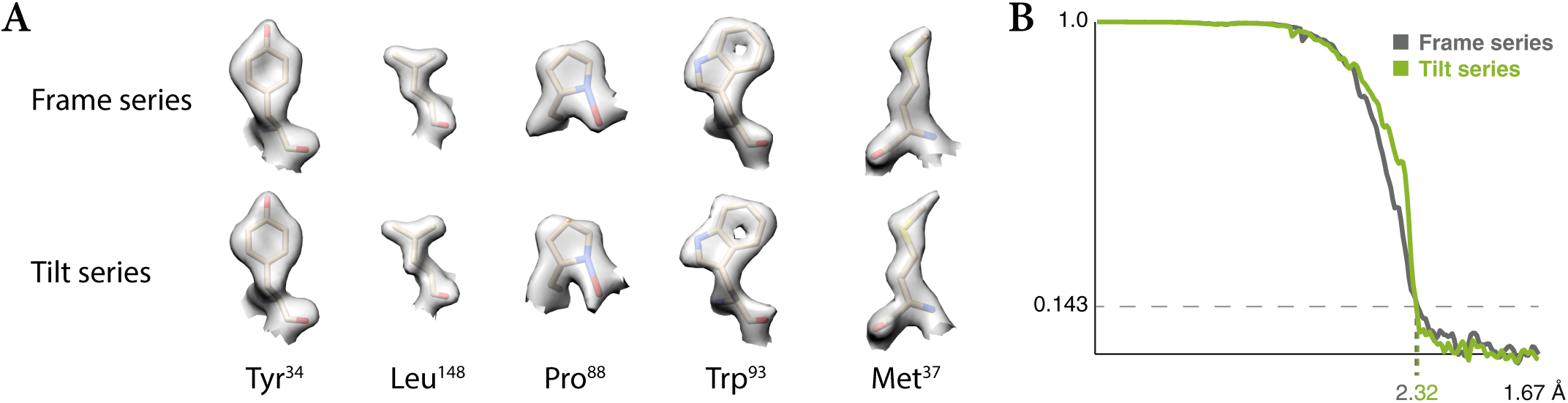
*M* achieves similar resolution for frame series and tilt series data. Given equal amounts of frame series and tilt series data of similar quality, *M* can achieve identical resolution, closing the gap previously assumed between both data types. This is exemplified here with the use of apoferritin data collected in both ways on the same grid. (a) Representative side chain densities observed in the frame series and tilt series maps. (b) Comparison between the global FSC curves for each map.

### Map denoising and local resolution

Instead of using a traditional Fourier Shell Correlation (FSC)-based approach for local resolution estimation^27^, *M* trains a deep learning-based denoiser model using a species’ half-maps to filter them to local resolution for the next refinement iteration. Because the noise estimation and filtering are done for each half-map independently, no common artifacts are introduced that could be amplified over subsequent refinement iterations. Even without local resolution filtering, artifacts may be introduced and amplified in regions of significantly lower resolution.

To assess the benefits of *M*’s denoising, we processed the EMPIAR-10288 data set containing the membrane protein cannabinoid receptor 1-G^28^. The 3.0 Å map published with the original study (EMD-0339) showed overfitting artifacts in the lipid bilayer (Fig. S3a). Processing the data with Warp, RELION and *M* led to only slightly improved resolution of 2.9 Å (Fig. S3b) using 149,308 particles (ca. 15% fewer than in the original study). However, the overfitting artifacts were absent in *M*’s final reconstruction (Fig. S3a). This is in line with improvements recently demonstrated using different approaches to local map filtering that do not rely on conventional FSC-based estimates^29, 30^.

### Comparison with RELION on atomic-resolution frame series data

To assess whether *M* can provide further improvements for frame series data processed with RELION 3.1, we refined a previously published^31^ apoferritin data set (EMPIAR-10248). The data were acquired on a novel JEOL microscope with a cold field emission gun to achieve an atomic resolution of 1.54 Å. Adding *M* to the pipeline improved the resolution to 1.35 Å, revealing densities for hydrogen atoms (Fig. S4). This shows that the image artifact model implemented in *M* can correct data at the highest end of the SPA resolution currently possible. At this high resolution, we were also able to assess the effect of Ewald sphere correction with the single side-band algorithm^25^. Applying it to the reconstruction alone, as would be possible in RELION 3.0, improved the resolution from 1.44 to 1.41 Å. Correcting the particle data and considering the sphere curvature during the multi-particle system refinement, improved the resolution further to 1.34 Å. Coupled with the demonstrated benefits of multi-species refinement and map denoising, this makes *M* a useful addition to the frame series SPA pipeline.

### Comparison with other tools for tilt series data refinement

To compare *M*’s reference-based tilt series alignment performance with the previously published EMAN2^15^ and emClarity^14^ packages, we reprocessed some of the data sets used in the respective publications (Fig. 5). EMAN2 reached a resolution of 8.4 Å on an *in vitro* 80S ribosome sample (EMPIAR-10064), improving significantly upon a previous 13 Å result^32^. For emClarity, a resolution of 8.6 Å was reported for the same data^14^. *M* improved the resolution to 5.7 Å and resulted in a map that clearly showed secondary structure elements and the helical groves of the RNA (Fig. 5a). We attribute a significant part of this improvement to *M*’s application of constraints between individual particle tilt images, which is not part of EMAN2.

**Figure 5.**
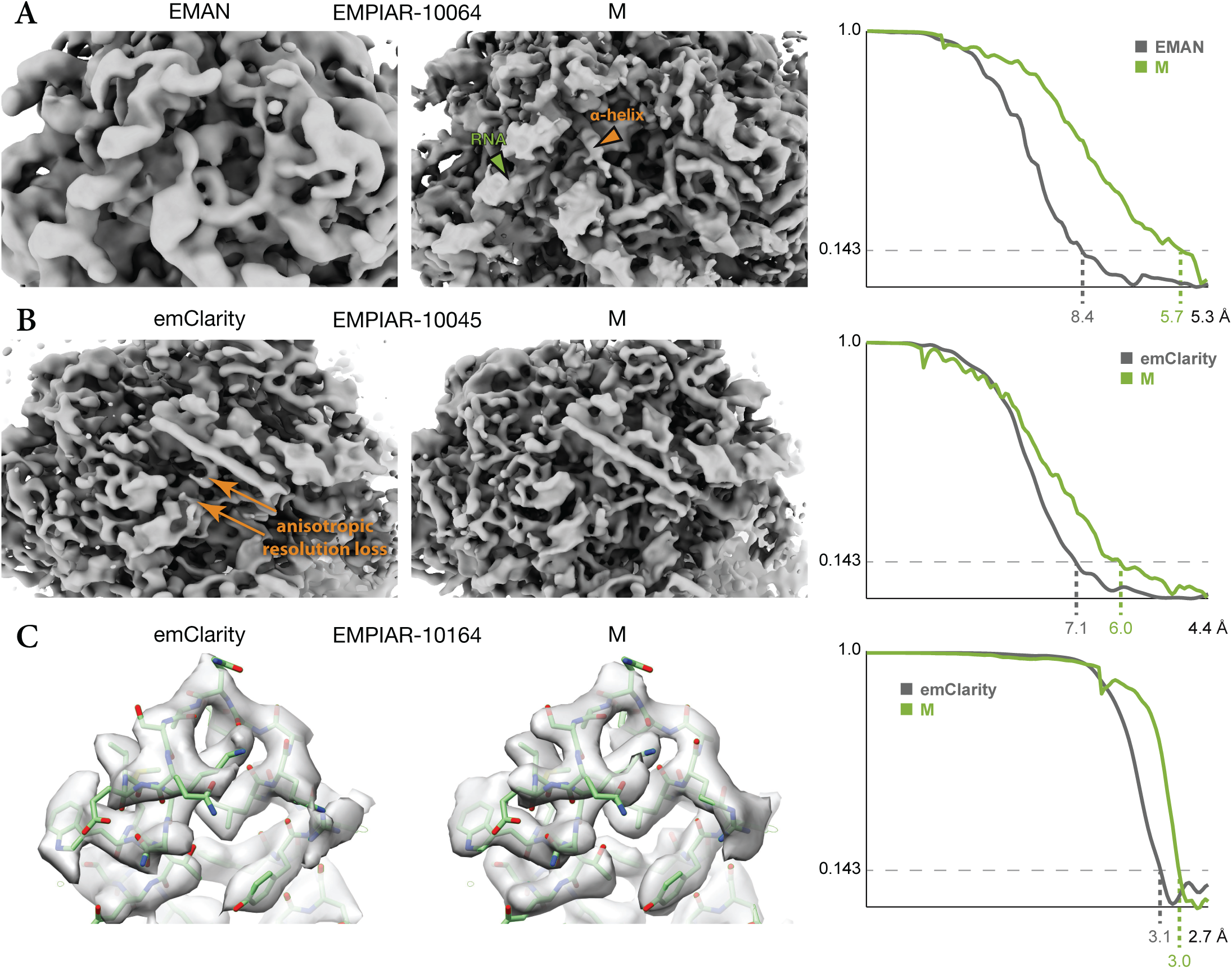
Comparison of maps obtained from published tilt series using *M* or other software. When applied to data used previously to test the EMAN^15^ and emClarity^14^ packages, *M* produces maps that compare favorably in terms of resolution and visual features. (a) 80S ribosome data from EMPIAR-10064 were used to benchmark the new tilt series processing in EMAN (EMD-0529). *M* achieved significantly higher resolution, accompanied by visibly better resolved features such as RNA (green arrow) and α-helices (orange arrow). (b) 80S ribosome data from EMPIAR-10045 were used to benchmark emClarity. The originally published map (EMD-8799, not shown) exhibited strong resolution anisotropy. A more recently updated map shown here^33^ still suffered from resolution anisotropy (“smearing” direction indicated by orange arrows). *M* achieved a slightly higher and more isotropic resolution, aiding the map’s interpretability. (c) HIV-1 capsid-SP1 data from EMPIAR-10164 were used to benchmark emClarity. Here, M achieved slightly higher resolution using ca. 30% of the particle number used by emClarity. Doubling the number of particles did not increase the resolution.

We further tested *M* on two data sets used in emClarity’s benchmarking. The emClarity software reached a resolution of 7.8 Å on purified 80S ribosomes (EMPIAR-10045) in the original publication^14^, and was later improved to 7.1 Å^33^, improving significantly upon a previous 12.9 Å result^20^. *M* improved the resolution to 6.0 Å, accompanied by improved resolution isotropy and map features (Fig. 5b). We attribute the improved resolution isotropy to *M*’s denoising-based map filtering approach that learns the optimal filtering empirically, whereas emClarity employs a FSC-based approach that may have to be tuned more conservatively to achieve the desired robustness to artifacts.

It was also reported that emClarity achieved a resolution of 3.1 Å on a thicker sample with a locally high particle density of isolated HIV-1 capsid-SP1 assemblies (EMPIAR-10164), improving upon previous 3.9 Å^34^ and 3.4 Å^35^ results. *M* improved the resolution to 3.0 Å, accompanied by local improvements in map quality (Fig. 5c). We attribute the slight improvement of overall resolution in both data sets to *M*’s more accurate deformation model and simultaneous optimization of all parameters, in contrast to emClarity’s separate steps for full image alignment (performed in IMOD^19^) and particle alignment (performed in emClarity). Our results show that *M* can improve over current methods, and achieve higher resolution with various tilt series data sets.

### *M* enables the visualization of an antibiotic bound to 70S ribosomes at 3.**7 Å *in situ***

To assess *M*’s performance on *in situ* data in the strictest sense, i.e. in tilt series obtained from intact cells, we used a data set of chloramphenicol-treated *Mycomplasma pneumoniae*^36^. *M* was able to resolve the 70S ribosome at 3.7 Å (Fig. 6a,d) based on 24,202 particles from 65 tomograms. The obtained map exhibited a wide range of local resolution values (Fig. 6b,d). The large 50S ribosomal subunit dominates the alignment and had a higher average resolution than the small 30S subunit, with much of its core reaching the 3.4 Å Nyquist limit of the data. In contrast, processing these data with Warp and RELION alone led to a 10 Å-resolution map of the 70S ribosome (Fig. 6c). M’s result constitutes a dramatic improvement compared to the previously used Warp-RELION pipeline, leading to a striking increase in structural detail (Fig. 6e).

**Figure 6.**
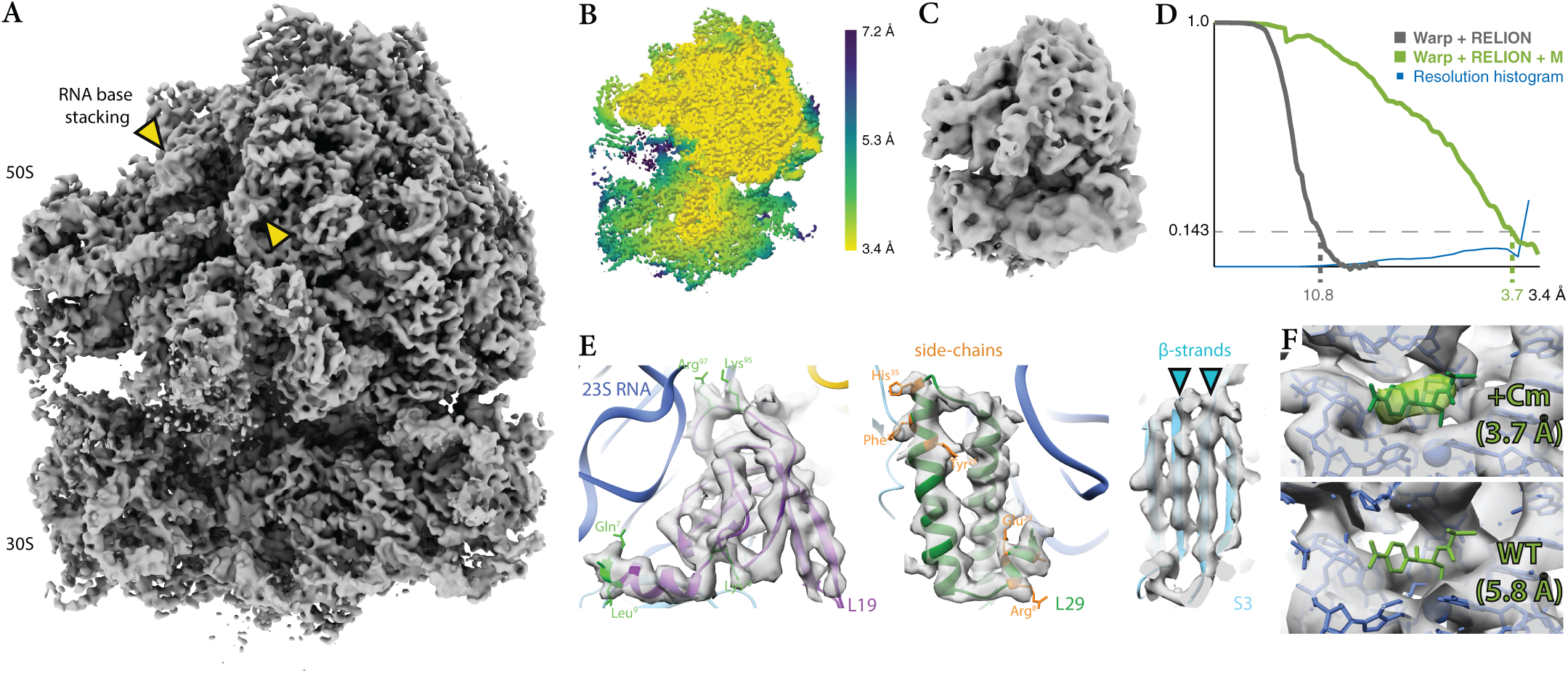
*M. pneumoniae* 70S ribosome-antibiotic map at 3.7 Å refined from *in situ* data with the new Warp–RELION–*M* pipeline. We applied the Warp-RELION-*M* pipeline to an *in situ* tilt series data set^36^. The achieved resolution reveals residue-level detail and a bound molecule of the antibiotic chloramphenicole (Cm). (a) Isosurface representation of the 3.7 Å resolution map. (b) Isosurface of the same map colored by local resolution. Despite stalling of the ribosome that is induced by antibiotic binding, residual ratcheting occurs that leads to higher resolution in the large 50S subunit, which dominates the alignment, and lower resolution in the small 30S subunit. (c) Isosurface of a 10.8 Å map derived from the same data set using only Warp and RELION shows the striking increase in detail after refinement with *M*. (d) Comparison between the FSC curves of the 3.7 Å and 10.8 Å map shows the increase in resolution achieved with *M*. The overlaid local resolution histogram of the 3.7 Å map shows that a significant portion of the map is resolved close to the data’s Nyquist limit of 3.4 Å. (e) High-resolution features, such as large amino acid side chains (orange arrow) and well-separated β-strands (green arrows), are resolved at a level expected for this resolution range. (f) Atomic model of a Cm-bound 70S ribosome (PDB-4v7w) fitted into the 3.7 Å map (top) shows correspondence of map density (light green) to the Cm molecule (dark green). Fitting the same model into a 5.6 Å *in situ* map of an untreated 70S ribosome (EMD-10683, bottom) does not show any density for Cm, providing a positive control.

The map possesses features typical for this resolution range, such as amino acid side chain stubs and β-sheets with individually resolved β-strands (Fig. 6e). A rigid body fit of an *E. coli* 70S ribosome–chloramphenicol structure (PDB-4v7t) further revealed the presence of a density corresponding to the chloramphenicol molecule at its expected target site (Fig. 6f), marking the first direct visualization of a drug bound to its target inside a cell. The density was absent in a 5.6 Å 70S reconstruction from untreated *M. pneumoniae* cells produced from processing with the older 1.0.6 Warp/*M* versions^36^ (Fig. 6f). Therefore, tilt series data acquisition on an intact cellular specimen with a mean thickness of 150 nm, in combination with the multi-particle refinement introduced in *M*, can lead to residue-level resolution structures of macromolecules in their native biological context.

## DISCUSSION

Here we present *M*, a cryo-EM map refinement tool that uses a multi-particle approach to obtain improved map resolution. Our results confirm that treating cryo-EM frame and tilt series as multi-particle systems rather than sets of isolated particles, and integrating their reference-based refinement with particle alignment and CTF refinement leads to improved map resolution. We show that the new framework removes previous technical limitations of tilt series data processing, allowing to achieve resolution at par with state-of-the-art frame series results, provided similar amounts of data. Although *M*’s refinement is constrained based on multi-particle assumptions, its image formation and reconstruction algorithms, as well as RELION’s 3D classification, assume isolated particles. While this is rarely an issue for *in vitro* data, refinement of crowded *in situ* data stands to benefit from extending these algorithms as well. Future work on reconstruction and classification algorithms may address this shortcoming by modeling the multi-particle system explicitly to achieve higher resolution. More generally, *M*’s optimization framework is flexible and can be extended in the future.

*M*’s ability to resolve structures *in situ* at a resolution previously considered exclusive to *in vitro* samples demonstrates that structures can be visualized directly inside cells at high resolution to arrive at atomic models. However, the ribosome is an outlier in terms of size and abundance in cells. Whereas significantly smaller complexes may be solved to similar resolutions in principle, the number of such instances may be limited due to the scarcity and heterogeneity of many complexes, and the difficulty of localizing them. Unlike *in vitro*, proteins cannot be significantly concentrated *in situ* without perturbing the organism. The only way to overcome this is to collect more data. Although both sample preparation and tilt series acquisition are becoming more streamlined, collecting enough particles of a rare protein complex to reach high resolution pose an impractical task for a single researcher or facility.

To help overcome this limitation, *M* offers data pooling and distributed processing mechanisms. These are powerful tools that can be of critical importance to share *in situ* image data and explore their potential by the community. Otherwise, only a very small portion of the data will be analyzed by individual research teams. We show that including more particle species in a multi-particle refinement improves the resolution of all particle species involved. Thus, everyone would benefit from having more proteins identified and refined in their data.

In conclusion, *M* can be combined with the established programs Warp and RELION into a powerful, semi-automated pipeline for cryo-EM data processing that includes a comprehensive and transferable sub-tomogram analysis workflow. This workflow avoids the need for conversion of file formats and conventions between different software packages, as currently common in the not yet streamlined processing of tilt series^37^, enabling non-expert users to achieve state-of-the-art results. It proves to have the potential to achieve residue-level resolution maps of particles inside cells and to capture macromolecular machines in action within their native environment. Together with complementary approaches, it further establishes the foundation for the emerging field of *in situ* structural biology.

## MATERIALS AND METHODS

### Data management

*M* requires data sources initialized based on a Warp project folder. Beside a list of frame/tilt series items, it stores the deformation model to be refined. *M* saves the refined deformation model for each item in the same XML metadata files previously created by Warp. Due to a shared code base, Warp can use the updated model when calculating new frame series averages or tomographic reconstructions. Multiple data sources of either type can be combined in a single population to facilitate the sharing and pooling of valuable in situ data that can contribute to far more than one project, but do not contain enough data for any single project on their own. To account for minor pixel size miscalibrations between different microscopes, the pixel size can be refined alongside other parameters in *M*.

A species is initialized from the refinement results of RELION or other compatible software, taking the unfiltered half-maps, a mask, and the particle coordinates and poses (i.e. translations and rotations) as a starting point. The state of a species after each refinement iteration comprises the reconstructed half-maps, the weights of the trained denoising model, various filtered and sharpened maps, a denoised map, and a list of particle coordinates and poses with multiple temporal sampling points if desired. The particles reference their data source items by their data hash to avoid naming conflicts between different data sources.

To enable multiple users to collaborate and pool their results, *M* tracks precisely the chain of refinements and other operations on data. After each refinement iteration, a “commit” is generated to save the new state. Similar to version-control systems like Git^38^, the commit’s hash is based on the exact state of the system committed. The hash of each data source item is calculated from the raw data, the refined deformation and imaging models, and the hashes of all species used for their refinement. The hash of each species is calculated based on the half-maps, the weights of the denoising model, the particle coordinates and poses, and the hashes of all data source items contributing information. The hashes can be used to verify a graph representing all steps that led to a particular state of a data source or species. Similar to the “pull request” mechanism in Git, species can be added to a population taking into account potential physical collisions with existing particles. This enables the maintenance of a centralized population repository from which multiple users can obtain prealigned data sources, identify new particle species or reclassify existing particles into more states, and contribute the results back to the repository.

### Deformation model

For frame series data, deformation of the multi-particle system is modeled in the XY plane only, with a pyramid of cubic spline grids^12^ *G*_*F, j*_ (*δ,i*) (where *j* is the index within the pyramid, *δ* is the spatial interpolation coordinate, and *i* is the temporal interpolation coordinate) going from high temporal/low spatial to low temporal/high spatial resolution. This accounts for the fast-changing, global stage movement, and the slowly-developing, local BIM. Furthermore, translation and rotation of individual particles as a function of exposure can be modeled with 2–3 control points depending on the particle size and overall exposure.

The model for tilt series data is more complex, owing to the higher potential for perturbations in the system between individual tilt exposures. As the mechanical rotation of the microscope stage and the estimated orientation of the tilt axis are imperfect, the assumed stage orientation can be randomly off in every tilt. *M* thus refines an independent set of stage rotation angle corrections *ω*_*i*_ for every tilt *i*. These corrections only affect the particle orientations to avoid redundancy, as the induced changes in the projected particle positions can be fully modeled by a deformation grid that must already be employed for other purposes.

Similarly, stage translation varies randomly between individual tilts. BIM patterns can be very different across adjacent tilt images as additional exposures are taken for focusing and tracking in-between. Particle positions can further deviate due to other imaging artifacts, such as wrongly calibrated magnification anisotropy^39^. *M* employs an “image warp” grid of cubic splines *GTI* with a spatial resolution of 3–5 in X and Y and per-tilt temporal resolution to model these geometric displacements in image space collectively. Furthermore, *in vitro* and *in situ* sample types for which tilt series are commonly used contain multiple overlapping layers of particles. Some deformations of densely filled volumes, such as shearing, or bending in the Z dimension when viewed at a high tilt angle, cannot be modeled accurately by XY translations in image space. *M* employs an additional “volume warp” grid *G*_*TV*_, implemented as a 4D grid of control points with quadrilinear interpolation between them that is anchored in volume space rather than image space. Hence it rotates with the sample and can model slow, continuous deformation that affects the particles’ projected positions in image space. As with frame series data, per-particle translation and rotation as a function of exposure is also modeled for tilt series.

Finally, a single tilt image exposure is usually fractionated in multiple frames, making it a tilt movie. At 1–3 e^-^/ Å^2^, the exposure in a single tilt movie is usually short, but still requires additional modeling to compensate motion. *M* parametrizes the XY translation as a combination of a grid with no spatial and perframe temporal resolution, and a grid with a spatial resolution of 3×3 and a temporal resolution of 3. Stage and particle orientations are assumed to remain constant throughout a tilt movie. Overall, the number of parameters for tilt series is larger than for frame series, requiring a higher particle density to achieve equivalent accuracy.

### Imaging model

The ability to model imaging conditions such as defocus, astigmatism, magnification or higher-order aberrations is equally important for obtaining high-resolution reconstructions. Frame and tilt series offer different advantages for refining some of these parameters.

For particles in frame series data, the Z coordinate and thus the relative offset from the global defocus of the micrograph is unknown. Although local defocus estimation based on amplitude spectrum fitting has been shown to increase resolution^12^, reference-based refinement of per-particle defocus can lead to a further increase in resolution^17^. *M* refines per-particle defocus and a per-series astigmatism for frame series, assuming constant values throughout the series.

Tilt series, on the other hand, provide accurate Z coordinates for all particles. However, the initial amplitude spectra-based global defocus estimates for each tilt have lower accuracy due to very short exposures, and cannot be assumed to remain constant throughout the series due to stage movement and refocusing. Furthermore, these estimates can be biased by contrast-rich objects that are not the particles of interest, such as a carbon film below or above the particles, or the platinum coating layer for FIB-thinned samples^40^. The astigmatism can also change between tilts due to fluctuating electron optics. *M* refines per-tilt defocus and astigmatism for tilt series, and calculates per-particle tilt CTFs based on these values and their Z coordinates. Particles in tilt series can potentially have more accurate defocus values because the number of parameters that can be fitted scales with the number of tilts or particles for tilt or frame series, respectively. In many cases the number of tilts will be significantly lower than the number of particles.

In both frame and tilt series, *M* also models per-series anisotropic magnification and higher-order optical aberrations. Refinement of a global set of Zernike polynomials representing the aberrations based on a 2D phase residual image calculated from all particles in a data set has been shown to improve the resolution significantly for slightly misaligned microscopes^41^. Within individual tilt series, too, beam tilt can vary as it is applied to compensate stage misalignments during tracking. Unfortunately, the signal in individual tilts is insufficient for accurate beam tilt estimation, and such an option is not implemented in *M*.

### Optimization procedure

*M* seeks to maximize the following target function *M*, which is essentially a weighted, normalized cross-correlation between all particle images and the corresponding reference projections:

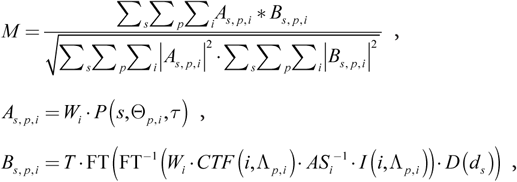

where *s* is a particle species, *p* is a particle of that species, and *i* is the index of a frame or tilt in a series; * denotes the dot product between two complex vectors, where the complex numbers are treated as pairs of scalars; |… | denotes the L_2_ norm; *W* is the anisotropic exposure- and tilt angle-dependent amplitude weighting of frame or tilt *i*; *P* is a projection operator in Fourier space sampling a central slice of the volume of species *s* at orientation Θ, taking into account the anisotropic scaling *τ*, bent to account for the Ewald sphere curvature determined by the species’ diameter; ·denotes scalar multiplication; *T* is the complex-valued beam tilt compensation; FT denotes the discrete Fourier transform; *CTF* is the real-valued CTF taking into account the defocus at position Λand the astigmatism in frame or tilt *i*; *AS* is the real-valued, rotational average over the amplitude spectra of all particle images of all species extracted from tilt *i* or the average of all aligned frames, used for spectrum whitening, scaled and cropped to the respective species size and resolution; *I* is the FT of a particle image extracted from frame or tilt *i* at position *δ*, cropped to the respective species resolution; *D* is a soft circular mask with particle diameter *d*.

Similar target functions in previous literature used *P. CTF* to model the contents of *I* ^16, 18^. However, in *M*’s implementation *I* is pre-multiplied by *CTF* to avoid CTF aliasing despite using small particle windows. This change does not affect the numerator part of *M* due to the associativity of complex number multiplication; its impact on the denominator part of *M* does not affect the achieved resolution in any way. It also avoids the additional memory footprint of storing pre-calculated CTFs, or the computational overhead of calculating them on-the-fly.

*M* can consider the Ewald sphere curvature during refinement if this is made necessary by a large species and/or high resolution^42^. In this case 2 copies of *CTF* ·*I* are prepared using the single side-band algorithm^25^: *CTF*_*P*_ ·*I* and *CTF*_*Q*_ ·*I*. To calculate the cost function, one is correlated with a bent central slice *P*, and the other with a central slice bent in the opposite direction. The resulting cost functions *M*_*P*_ and *M*_*Q*_ are then added. As with previous implementations^17^, the absolute handedness for the correction must be provided by the user.

For frame series, the position and orientation of particle *p* in frame *i* are calculated as:

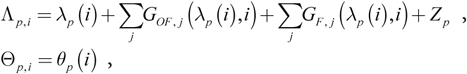

where *λ* is the value of the refined particle position trajectory interpolated at the accumulated exposure of frame *i*; *G*_*OF*_ is a deformation grid pyramid produced by Warp’s original refer-ence-free alignment that is not altered in *M* refinement; *G*_*F*_ is a deformation grid pyramid that is refined in *M*; *Z* is the refined defocus value of particle *p* that is added as the Z coordinate to its position; *θ* is the value of the refined particle orientation trajectory interpolated at the accumulated exposure of frame *i*.

For tilt series, the position and orientation of particle *p* in tilt *i* are calculated as:

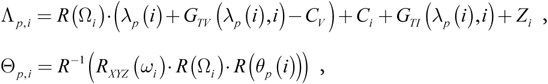

where *R* and *R*_*XYZ*_ construct a rotation matrix based on a set of Euler and XYZ angles, respectively, and *R*^-1^ calculates a set of Euler angles based on a rotation matrix; *C*_*V*_ is the center of the volume in which the multi-particle system is anchored, and *C*_*i*_ is the center of the full tilt image; *Z*_*i*_ is the refined defocus value of tilt *i* that is added to the Z coordinate of the transformed particle position; Ωis the stage orientation determined in the initial, reference-free tilt series alignment that is not altered in *M* refinement; ·denotes matrix multiplication here.

For frames in tilt movie *i*, the position of particle *p* in frame *k* is calculated as:

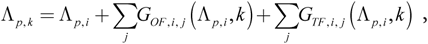

where *G*_*OF*_ is the deformation grid pyramid produced by Warp’s original reference-free alignment of the tilt movie that is not altered in *M* refinement.

Due to the very large number of parameters, *M* employs L-BFGS^22^ to perform almost all of the optimization. Only the initial defocus search is done exhaustively over a limited range to avoid getting trapped in a local optimum because of the quickly oscillating nature of the CTF. Every L-BFGS search iteration requires the calculation of a partial derivative of the target function with respect to each optimizable parameter. Reevaluating *M* twice per parameter to compute the gradient with the central differences numerical scheme would be very computationally expensive. Like Warp, *M* takes a computational shortcut for most of the parameters.

Before optimization starts, *M* calculates the partial derivatives of the X and Y components of all Λ_*p, i*_ with respect to all warping grid parameters and all control points of a particle’s position trajectory that affect them. Similarly, the partial derivatives of the individual Euler angle components of all Θ_*p, i*_ with respect to all stage angle correction parameters and all control points of a particle’s orientation trajectory are calculated. As each parameter influences only a small fraction of particle frames or tilts, most of the derivatives are 0. They are excluded from the precalculated lists to avoid unnecessary computation. Then, during optimization, once per search iteration, the partial derivative of 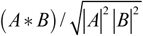 for each particle frame or tilt is calculated with respect to X, Y and the Euler angles. This amounts to evaluating *M* 10 times. A useful approximation for the derivative for each parameter *η* can then be calculated as follows:

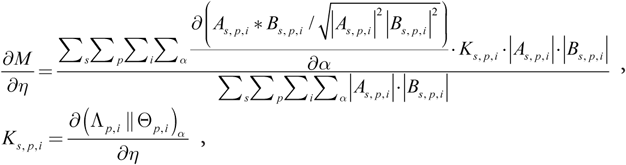

where *α*∈{*x, y,ϕ,ϑ,ψ*}, i. e. one of the translation axes or Euler angles; ‖ denotes the concatenation of two tuples; (…)_*α*_ denotes the selection of component *α* from a tuple.

The deformation parameters make up the bulk of all parameters. Parameters such as absolute magnification and beam tilt do not benefit from the same shortcut and their derivatives must be calculated independently with the central differences scheme. The CTF-related parameters are few, but the calculation of their derivatives is especially expensive because it requires the particles to be reextracted at an aliasing-free size, pre-multiplied by the altered CTF, and cropped to refinement size – all involving expensive FT steps. *M* calculates the values of *M* by adding up the results from small batches of particles. This allows the cost of the first FT at aliasing-free size to be amortized over all optimizable CTF parameters, as its result is reused for all subsequent calculations. The gradients for all per-particle or per-tilt defocus and astigmatism parameters can all be calculated in the same pass as each of them affects only one particle or tilt.

If defocus is to be optimized, an iterative grid search can be executed before the L-BFGS optimization starts. The search runs for 5 iterations. For the first iteration, a range of ±300 nm around the current values is sampled in 10 nm steps. For each subsequent iteration, the search step is halved, and a range of ± the new search step around the 2 best values for each particle or tilt from the previous iteration is sampled.

### Memory footprint considerations

Traditional SPA refinement treats every particle as an isolated entity, thus requiring no more than one particle to be held in memory at any given time if parallelization is not considered. A multi-particle approach, however, needs to rapidly evaluate the state of the entire multi-particle system during refinement. The particle frame/tilt series need to be stored in memory because re-extracting and reprocessing them for every evaluation would be too inefficient. While an *in vitro* sample usually contains a single layer of proteins with up to 1000–2000 particles in a field of view, a densely packed *in situ* volume has the potential to contribute tens of thousands of particles to refinement if enough species can be identified. The image size is selected to be twice the particle diameter to account for signal delocalization and interpolation artifacts, leading to significant overlap even in the single-layer case. At high refinement resolution, the memory requirements of all extracted particle frame/tilt series in a system can vastly exceed those of the original data, rising to tens or even hundreds of gigabytes.

Although *M* uses GPUs for acceleration wherever possible, currently available consumer-level cards offer up to 12 GB, which would be insufficient in many cases. Therefore, the extracted particle frame/tilt series are held in “pinned” (i.e. page-locked) CPU memory where they can be transparently accessed by the GPU. Despite the low bandwidth of CPU–GPU memory transfers, the GPU does not experience a significant performance penalty when correlating them to reference projections. This is because the particle data accesses are sequential and highly coalesced, whereas the creation of reference projections on-the-fly accesses the GPU memory randomly, creating significant overhead. As faster CPU–GPU interfaces are being developed, the penalty should become more negligible in the future.

Still, memory requirements can become too high even for CPU memory. To reduce the footprint, *M* exploits the varying information content of frames/tilts over the course of a series. As sample damage from radiation is accounted for by applying a Gaussian (“B-factor”) weighting function in Fourier space^7, 17^, the contribution of higher-frequency components becomes negligible at high exposure. *M* crops extracted particle images in Fourier space to a resolution that corresponds to the weighting function value falling below 0.25, resulting in considerable space savings once high resolution is reached. Assuming an increase in the weighting B-factor of 4 Å ^2^ per 1 e^-^/ Å ^2^ of accumulated exposure, the maximum useful frequency at exposure *d* is 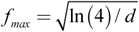, and the image size *m* scales with a factor of min(1, *f*_*max*_ / *f*_*refine*_). Thus, the upper bound for memory consumption in case of low refinement resolution and/or low overall exposure is *O*(*m*^2^ *d*), while the lower bound is Ω(*m*^2^ln (*d*)) in case of high refinement resolution and/or high overall exposure.

### Avoiding CTF aliasing

Cryo-EM data of thin biological specimens are usually acquired at defocus to achieve phase contrast. In the absence of a phase plate device, and often in the case of *in situ* tomography, defocus values can exceed 4 µm to enable better visual interpretation of the raw data. Higher defocus results in stronger delocalization of the signal in real space, as reflected by faster oscillations of the CTF in Fourier space. As the CTF oscillates between -1 and 1, combining signals with different defoci would result in an average value of 0 at higher spatial frequencies. Thus, a phase shift of *π*must be applied to frequency components modulated by negative CTF values prior to averaging. Furthermore, it is desirable to compute the reconstruction as a weighted average, using the CTF for the weighting. Multiplying the FT of a particle image by the corresponding real-valued CTF achieves both goals.

Current SPA packages advise the user to select the particle box size as 1.5–2 the particle diameter to account for Fourier-space interpolation artifacts, not considering the image defocus. When an image is cropped around a particle, the Fourier-space modulation pattern becomes band-limited to the new window size. If CTF oscillations are too fast to be resolved, the band-limited values for the amplitudes of the corresponding frequency components will converge to 0. Even worse, the analytical 2D CTF model used in refinement and reconstruction is not band-limited, and contains solely aliasing artifacts past the Fourier-space Nyquist frequency instead of converging to 0. This can put a hard limit on the achievable resolution for small particles and those acquired at high defocus that is independent of the actual data quality.

This problem can be mitigated by selecting a box size large enough to avoid CTF aliasing at the highest defocus value in a data set. However, the required size *m* can exceed 1000 px at high resolution or defocus, significantly slowing down refinement algorithms whose complexity and memory footprint are *O* (*m*^2^) and *O*(*m*^3^), respectively. This increase can be entirely avoided by pre-multiplying particle images by the CTF at an aliasing-free size, and cropping them to a smaller size for refinement or reconstruction. As the modulation pattern is CTF^2^ after pre-multiplication, the band-limited oscillations will converge to 0.5 instead of 0. The 2D CTF model used in refinement and reconstruction must be similarly band-limited to match the data. As *M* operates on all particles of an entire frame/tilt series at a time and extracts the particle images on-the-fly, such considerations are made automatically for the currently needed resolution.

The minimum box size needed for CTF correction at a given resolution is dictated by the maximum oscillation rate of the CTF within the available spatial frequency range. This is not necessarily the oscillation rate at the highest spatial frequency as *φ*is not a monotonic function: A combination of low underfocus and high C_s_ will cause the oscillations to slow down significantly and accelerate again at higher spatial frequencies. The oscillation rate can be calculated as the first derivative of *φ*. In practice, it is easier to evaluate *dφ*/ *dk* numerically within the relevant range of spatial frequencies to find its maximum absolute value. To fully resolve the oscillation, one period must be rasterized onto at least 2 pixels, i.e. the window size must be chosen such that max (*dφ*/ *dk*) = 2*π*/ 2 *px*. While this guarantees a fully resolved CTF in 1D, a CTF rasterized on a Cartesian 2D grid has an anisotropic sampling rate. At its lowest, *i*.*e*. along the diagonals, it requires 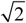 the sampling rate of the 1D case.

Before particle extraction, the size padding factor at which the images will be pre-multiplied by the CTF has to be determined, taking into consideration the maximum defocus value expected in a frame/tilt series, and the expected maximum resolution. During refinement, the latter is set to the refinement resolution. For the final reconstruction, it is set to 1.25x the current global resolution. Particles are extracted using the calculated minimum box size (or twice the particle diameter in case that value is larger), and pre-multiplied by the CTF in Fourier space. Then the inverse FT (IFT) is applied, the particles are cropped to the refinement or reconstruction size in real space, and transformed back to Fourier space for refinement. The band-limited CTF^2^ model is prepared by simulating the function at the same aliasing-free size in Fourier space, cropping its IFT in real space, and taking the real components of the result’s FT.

### Data-driven weighting

To account for radiation damage as a function of accumulated exposure, or increasing sample thickness as a function of the stage orientation, several heuristics and empirical approaches have been proposed^7, 17, 20^. By default, *M* adopts the heuristic introduced in RELION 1.4^20^. The B-factor is increased by 4 Å ^2^ per 1 e^-^/Å^2^ of exposure, and each tilt is weighted as cos *ϑ*. Once high resolution is reached, the weights can be estimated empirically using a reference correlation-based approach similar to the one introduced in RELION 3.0^17^.

In a departure from RELION’s scheme, the normalized correlation (NC) is calculated between particle images and reference projections at the end of a refinement iteration are not combined across the entire data set. It is kept as a 2D image to enable the fitting of anisotropic weights rather than averaging rotationally. The correlation data can then be recombined in different ways to calculate different kinds of weights. Furthermore, because *M* supports the refinement of multiple species with different resolution, the per-species correlation vectors for each frame or tilt need to be combined. This is done by weighting each one by the FSC calculated between the half-maps of the respective species. This produces a set of vectors *NC*_*d, i, k*_, where *d* is the series, *i* is the frame or tilt, and, optionally, *k* is the tilt movie frame.

The procedure then iteratively calculates 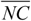 as:

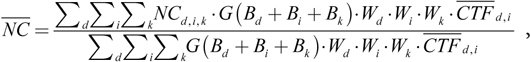

and optimizes the weighting parameters to minimize the following cost function:

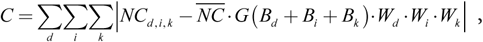

where ·denotes scalar multiplication; *G* is an anisotropic 2D Gaussian B-factor weighting function; *B* is a vector describing the B-factor along the X and Y axes, and their rotation; *W* is a scalar weight; 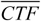 is the weighted average of all particle CTFs in one frame or tilt. The B-factors in each group are constrained such that the highest value in a group is set to 0.

In this default formulation, the weighting scheme allows to assign separate weights not only to individual frames/tilts, but also to weight the contribution of an entire series. For data with high particle density this scheme can be extended to assign different weights to frames/tilts of each individual series. Anisotropic B-factors improve the weighting of frames with significant intra-frame motion (Fig. S5). Combined with per-series, perframe weighting, such granularity allows to rescue more information from the first few frames of an exposure if parts of them are less affected by BIM.

### Map reconstruction

Previous refinement packages took two different approaches to map reconstruction from frame and tilt series data. For frame series, weighted averages were prepared either directly from the initial, reference-free alignments, or based on a “polishing” procedure^17^. These 2D averages were then weighted based on a 2D CTF model and a spectral signal-to-noise ratio (SSNR) term^18^, and back-projected to obtain the reconstruction. For tilt series, the algorithms operated on intermediate per-particle 3D reconstructions (‘sub-tomograms’) with fixed translational and rotational offsets between individual tilt images. These 3D sub-tomograms were then weighted based on a 3D CTF model^20^ and an SSNR term, and back-projected to obtain the reconstruction.

*M* seeks to unify the handling of both types of data and uses the original, non-interpolated 2D data at every step, including reconstruction. For tilt series, this approach avoids any artifacts from intermediate interpolation and reconstruction steps. For frame series, the requirement for identical orientation of all particle frames no longer exists as they are not averaged in 2D, enabling the modeling of particle orientation as a function of exposure. Only for individual tilt movie frames a shortcut is taken to save memory and computation, and they are pre-averaged in 2D using the approach described for Warp^12^ after a separate multi-particle refinement of the respective tilt movie.

Thus, for the reconstruction, individual particle frames or tilts are weighted by an exposure-dependent function to account for radiation damage, and an aliasing-free 2D CTF model (see previous section) that incorporates the exact defocus and astigmatism values for that position and frame/tilt. The weighted data are then back-projected through Fourier space summation, accounting for Ewald sphere curvature. The reconstruction is finalized by dividing the summed data component by the summed weights component^18^.

### Map denoising

Reconstructions of biological specimens derived from cryo-EM data rarely have homogeneous resolution throughout all parts of the macromolecule. Using a map filtered to its global resolution for particle alignment can have detrimental effects. Poorly resolved regions, such as floppy protein domains or the lipid bi-layer around transmembrane domains, will make the alignment worse by adding noise to reference projections below the refinement resolution. In the case of fully independent half-maps^43^, the noise patterns that the particles will be aligned against are independent, and amplifying them over several iterations only has the potential of making the resolution worse. In the case of refinement with merged half-maps^16^, where overfitting is avoided by limiting the refinement resolution, the poorly resolved regions may be well below that limit, leading to a common, overfitted noise pattern in both half-maps.

Past attempts at filtering maps based on local resolution estimates for refinement^44, 45^ applied FSC-based approaches^27^ to estimate the local resolution and performed the filtering in the Fourier domain. As only one set of estimates can be made based on one pair of half-maps, any spurious patterns in the estimated values will be introduced into both half-maps when the filtering is performed. The locality and accuracy of the estimates depends on the window size^27^. A smaller window increases locality at the expense of accuracy. Once introduced, the noise pattern can become amplified over multiple iterations, leading to overestimated local resolution and phantom features that can be misinterpreted. More advanced regularization schemes have been proposed^29, 30^ since to deal with this problem.

*M* implements a new approach to map filtering that uses neural network-based denoising. After the recently proposed noise2noise training principle^46^ has been successfully applied to micrograph^12^ and tomogram^12, 47^ denoising, half-map reconstructions provide another obvious case of two independently noisy observations of the same signal. We find that a denoiser trained on one pair of half-maps not only matches closely the result of conventional global resolution filtering when applied to maps with homogeneous resolution, but also provides locally smooth, artifact-free local resolution filtering. As such models can train on and denoise sets of micrographs or tomograms with different defocus values and thus different noise models, they can also recognize and adapt to different noise levels within the same reconstruction. In another important departure from FSC-based methods, the denoising step is applied to the half-maps independently and the denoiser sees only one of them at a time. Thus, even if some spurious pattern is introduced as part of the denoising, it is independent between the half-maps.

The neural network architecture is identical to the one used for tomogram denoising in Warp. A separate denoising model is maintained for every species, and trained only on the respective pair of half-maps. The model is initialized with random values and trained for 800 iterations upon the creation of a new species. It is later retrained for another 800 iterations after every refinement. Spectrum whitening is applied to the maps before training to restore high-frequency amplitudes^16^, similar to B-factor-based sharpening^48^. During training, 64^3^ px volumes are extracted from both maps at the same random position and orientation, and presented to the network as input and output in mini-batches of 3. The random orientations make sure the network learns the noise model rather than merely learning the average map. The learning rate for the Adam optimizer is exponentially decreased from 10^−3^ to 10^−5^ throughout the training. For the denoising of each half-map, the map is partitioned in 64^3^ px windows overlapping by 24 px, denoised, and the results from each window are inserted into the output volume. Regardless of regions with above-average resolution being potentially present, the refinement resolution is set conservatively to the global map resolution. In addition to the two half-maps for refinement, a denoised average map is also prepared by applying the same denoising model to the average of the spectrum-whitened half-maps.

### Assessment of map denoising

Frame series data were downloaded for the EMPIAR-10288 entry (Fig. S3). Frame alignment and local CTF estimation were performed in Warp with a spatial resolution of 5×5. 1,033,994 particles were picked with a retrained BoxNet model in Warp and exported at 1.5 Å/px. 2D classification, 3D classification and refinement were performed in RELION using EMD-0339 as the initial reference. 149,328 particles corresponding to the best 3D class were imported in *M*. The particle poses were given a temporal resolution of 2, the deformation grid resolution was set to 2×2, and refinement of all parameters was performed for 5 iterations. Data-driven weight estimation was performed to assign unique weights to every frame index.

### Acquisition of apoferritin benchmark data

To compare the resolution achievable with frame and tilt series data and assess individual algorithms implemented in *M*, we acquired two data sets of human heavy-chain apoferritin: AF-f (frame series) and AF-t (tilt series). To make sure that any observed differences came from data type and processing strategies rather than local variance in sample quality, neighboring holes within the same grid square were used for both data sets.

The apoferritin plasmid and purification protocol were kindly provided by Louise Fairall and Christos Savva from the Midlands Regional Cryo-EM Facility, University of Leicester. In brief, GST-tagged apoferritin was overexpressed in *E. coli*, captured on Gluthatione-sepharose beads after cell lysis, cleaved off the resin by TEV protease and purified to homogeneity by size exclusion chromatography in 50 mM Tris-HCl pH 7.5, 100 mM NaCl and 0.5 mM TCEP.

3 ml of apoferritin at 3.8 mg/ml were applied to freshly glow discharged R 1.2/1.3 holey carbon grids (Quantifoil) at 4°C and 100% relative humidity followed by plunge-freezing in liquid ethane using a Vitrobot Mark IV (Thermo Fisher Scientific). The sample concentration resulted in a dense, single-layered hole coverage. Data were collected on a Titan Krios TEM (Thermo Fisher Scientific) operated at 300 kV and a magnification resulting in a calibrated pixel size of 0.834 Å. The energy filter (Gatan) was operated in zero loss mode with a slit width of 20 eV. The K3 direct electron detector (Gatan) was operated in counting mode with a freshly acquired reference for gain correction. The exposure rate was adjusted to 20 e^-^/px/s. SerialEM^49^ was used for frame and tilt series acquisition.

Positions for both data sets were selected to be distributed evenly over the same grid area to maximize the similarity in ice thickness and particle density. For AF-f, 150 frame series were collected with a total series exposure of 32 e^-^/Å^2^, fractionated in 40 frames. For AF-t, 135 tilt series ranging from -40 to +40 degrees were collected in a grouped dose-symmetric scheme^50^ with a group size of 2 and in 2 degree steps. Each tilt was exposed to 2.7 e^-^/Å^2^, fractionated in 3 frames.

### Comparison between frame and tilt series performance

Using data set AF-f, frame series alignment and local CTF estimation were performed in Warp with a spatial resolution of 8×5, owing to the rectangular format of the K3 chip. 22,122 particles were picked with a retrained BoxNet model in Warp and exported at full resolution in 512 px boxes. Global 3D refinement with octahedral symmetry was performed in RELION 3.0. The results were imported in *M*. The particle poses were given a temporal resolution of 3, the deformation grid resolution was set to 6×4, and refinement of all parameters was performed for 5 iterations. Data-driven weight estimation was performed to assign unique weights to every series and frame index.

Using data set AF-t, tilt movie frame alignment was performed in Warp using a model without spatial resolution. Initial tilt series alignment was performed in IMOD using patch tracking on 6x binned images with default settings. Tilt series CTF estimation was performed in Warp. 18,991 particles were picked using Warp’s 3D template matching in full tomograms reconstructed at 10 Å/px. Sub-tomograms and 3D CTF volumes were exported at 2 Å/px using 140 px boxes. Global 3D refinement with octahedral symmetry was performed in RELION 3.0. The results were imported in *M*. The particle poses were given a temporal resolution of 3, the image warp grid resolution was set to 6×4×41, and refinement of all parameters was performed for 5 iterations, including tilt movie frame alignment in the last 2 iterations. Data-driven weight estimation was performed to assign unique weights to every series and tilt index.

### Assessment of multi-species refinement

Particles from each frame series of the AF-f data set were split in 5% and 95% sub-populations, resulting in species with 3,710 and 70,497 particles, respectively. Frame alignments and particle poses previously obtained from Warp and RELION were reused. In the first scenario, the 5% species was refined alone. In the second scenario, the 5% species was co-refined with the 95% species. Both species were assumed to be structurally independent and did not contribute particles to each other’s reconstructions. For both tested scenarios, a 6×4 starting grid for the deformation was used, the resolution of all species was set to 4.0 Å and only one refinement iteration was performed in *M* to avoid possible benefits from the higher resolution the 95% species would reach after the first iteration.

### Comparison with RELION on atomic-resolution frame series data

Frame series data were downloaded for the EMPIAR-10248 entry and pre-processed in Warp. 109,437 particles were exported at 0.6 Å/px using 466 px boxes and refined in RELION. The resulting particle poses and half-maps were imported in *M* and refined for 5 iterations starting with a resolution of 3.0 Å in the first iteration. A starting grid of 4×4 was used for the deformation model. All CTF-related parameters were refined, including per-series beam tilt and a 3×3 grid model for local astigmatism. For the last 2 iterations, anisotropic per-series, per-frame B-factor weights were estimated. The original mask deposited with EMD-9865 was used to estimate the final resolution.

### Comparison with other tools for tilt series data refinement

Tilt series movie data were downloaded for the EMPIAR-10164 entry. Tilt movie frame alignment was performed in Warp using a model without spatial resolution. Initial tilt series alignment was performed in IMOD using gold fiducials automatically picked in Warp, on 6x binned images with default settings. Tilt series CTF estimation was performed in Warp. 130,658 particles were picked using Warp’s 3D template matching with a template derived from EMD-3782 in full tomograms reconstructed at 10 Å/px. Sub-tomograms and 3D CTF volumes were exported at 5 Å/px using 56 px boxes. Global 3D refinement with C6 symmetry was performed in RELION 3.0, and reached the 1/10 Å ^-1^ Nyquist frequency. The results were imported in *M*. The particle poses were given a temporal resolution of 3, the image warp and volume warp grid resolutions were set to 8×8×41 and 3×3×3×20, and refinement of all parameters was performed for 5 iterations, including tilt movie frame alignment in the last 2 iterations. Data-driven anisotropic weight estimation was performed to assign unique weights to every series, tilt index and tilt frame index.

Tilt series data were downloaded for the EMPIAR-10045 entry. Initial tilt series alignment was performed in IMOD using manually picked gold fiducials on 4x binned images with default settings. Tilt series CTF estimation was performed in Warp. 3,058 particles were picked using Warp’s 3D template matching in full tomograms reconstructed at 10 Å/px. Sub-tomograms and 3D CTF volumes were exported at 5.0 Å/px. Global 3D refinement reached a resolution of 13 Å. The results were imported in M. The particle poses were given a temporal resolution of 3, the image warp and volume warp grid resolutions were set to 8×8×41 and 4×4×2×20, respectively, and refinement of all parameters was performed for 5 iterations. Data-driven anisotropic weight estimation was performed to assign unique weights to every series and tilt index.

Tilt series data were downloaded for the EMPIAR-10045 entry. Initial tilt series alignment was performed in IMOD using manually picked gold fiducials on 4x binned images with default settings. Tilt series CTF estimation was performed in Warp. 3,566 particles were picked using Warp’s 3D template matching in full tomograms reconstructed at 10 Å/px. Sub-tomograms and 3D CTF volumes were exported at 5.0 Å/px. Global 3D refinement reached a resolution of 13 Å. The results were imported in M. The particle poses were given a temporal resolution of 3, the image warp and volume warp grid resolutions were set to 8×8×41 and 4×4×2×20, respectively, and refinement of all parameters was performed for 5 iterations. Data-driven anisotropic weight estimation was performed to assign unique weights to every series and tilt index.

### Acquisition and refinement of *M. pneumoniae in situ* tilt series data

Data previously used in another study^36^ were re-analyzed with the release version of *M*. As described there, *Mycoplasma pneumoniae* strain M129(ATCC 29342)cells were grown on 200 mesh gold grids coated with a holey carbon support (R 2/1, Quantifoil). Cells were cultivated at 37 °C in modified Hayflick medium: 14.7 g/L Difco PPLO (Becton Dickinson, USA), 20% (v/v) Gibco horse serum (New Zealand origin, Life Technologies, USA), 100 mM HEPES-Na (pH 7.4), 1% (w/w) glucose, 0.002% (w/w) phenol red and 1,000 U/mL freshly dissolved penicillin G. Chloramphenicol (Cm; Sigma-Aldrich, USA) was added 15 minutes prior to vitrification, at a final concentration of 0.5 mg/ ml. Grids were quickly washed with PBS buffer containing 10 nm protein A-conjugated gold beads (Aurion, Netherlands), blotted from the back side for 2 seconds, and plunged into mixed liquid ethane/propane at liquid N_2_ temperature with a manual plunger (Max Planck Institute of Biochemistry, Germany). The cryo-EM grids were stored in a sealed box in liquid N_2_ before usage.

Tilt series data were collected on a Titan Krios TEM operated at 300 kV (Thermo Fisher Scientific) equipped with a field-emission gun, a Gatan K2 Summit direct detector and a Quantum post-column energy filter (Gatan). Images were recorded in exposure-fractionation, counting mode using SerialEM 3.7.2. Tilt-series were acquired with a dose-symmetric scheme using dedicated scripts^51^ with the following settings: TEM in nano-probe mode, magnification 81,000 with a calibrated pixel size of 1.7 Å, energy filter in zero loss mode, defocus range 1.5 to 3.5 µm, tilt range -60° to 60° with 3° tilt increment and constant exposure per tilt, total exposure of 120 e^-^/Å^2^. In total, 65 tilt series were collected from Cm-treated cells.

Raw tilt movies were processed in Warp. *De novo* tilt series alignment was performed in IMOD using gold fiducials picked automatically with Warp’s BoxNet, and the results were imported in Warp, where the tilt series CTFs were estimated. Using full tomograms reconstructed at 10 Å/px, two tomograms were denoised using Warp’s Noise2Map tool to pick the ribosome particles manually. Using these coordinates, sub-tomograms were exported from Warp to RELION to obtain an initial reference. This reference was used to perform template matching in Warp at 10 Å/px. In addition, a 3D convolutional neural network was trained on the 2 manually picked tomograms to remove false positives (membranes, carbon hole edges etc.) from the template matching results. 24,202 particles were obtained this way. Sub-tomograms for all particles were exported from Warp to RELION and aligned against the previously refined low-resolution reference. No classification was performed. The results were imported in *M*. There, global movement and rotation, a 5×5×41 image-space warping grid, a 8×8×2×10 volume-space warping grid, as well as particle pose trajectories with 3 temporal sampling points were refined over 5 iterations. Starting with iteration 3, CTF parameters were also refined. At the beginning of iteration 4, reference-based tilt movie alignment was performed. Similar results for the final map (not shown here), demonstrating the flexibility of *M*, were obtained independently using on-the-fly raw tilt movies alignment in SerialEM^49^ and template matching in pyTOM^52^.

## ACKNOWLEDGEMENTS

The human H-chain apoferritin plasmid and purification protocol were kindly provided by the Loise Fairall, Christos Savva and the Protex facility of the University of Leicester. P.C. was supported by the Deutsche Forschungsgemeinschaft within SFB860 and SPP1935 and under Germany’s Excellence Strategy (EXC 2067/1-390729940), the European Research Council Advanced Investigator Grant TRANSREGULON (grant agreement No 693023), and the Volkswagen Foundation. J.M. acknowledges support from the EMBL and a European Research Council starting grant 3DCellPhase (760067).

## AUTHOR CONTRIBUTIONS

D.T. designed *M*’s architecture and all algorithms, and carried out all implementation and application. L.X. and J.M. provided tilt series data of chloramphenicol-treated *M. pneumoniae* cells, assisted in testing *M* and interpretation of the maps. D.T. and L.X. solved the Cm-bound ribosome structure. C.D. collected apoferritin frame series and tilt series for benchmarking, and analysed the frame series data using RELION. P.C. provided scientific environment, funding and additional interpretations and implications. D.T., P.C. and J.M. wrote the manuscript with input from all authors.

## CODE AVAILABILITY

M and Warp binaries, source code and user guide can be downloaded from https://github.com/cramerlab/warp and https://warpem.com.

## DATA AVAILABILITY

Coulomb potential maps for the *in situ* 70S ribosome, apoferritin frame and tilt series comparison, and EMPIAR-10248 reprocessing will be deposited in the Electron Microscopy Data Bank (EMDB). Raw *in vitro* data for the apoferritin frame and tilt series comparison, and the raw *in situ* data of Cm-treated *M. pneumoniae* cells will be deposited in the EMPIAR database.

---

**Figure S1.**
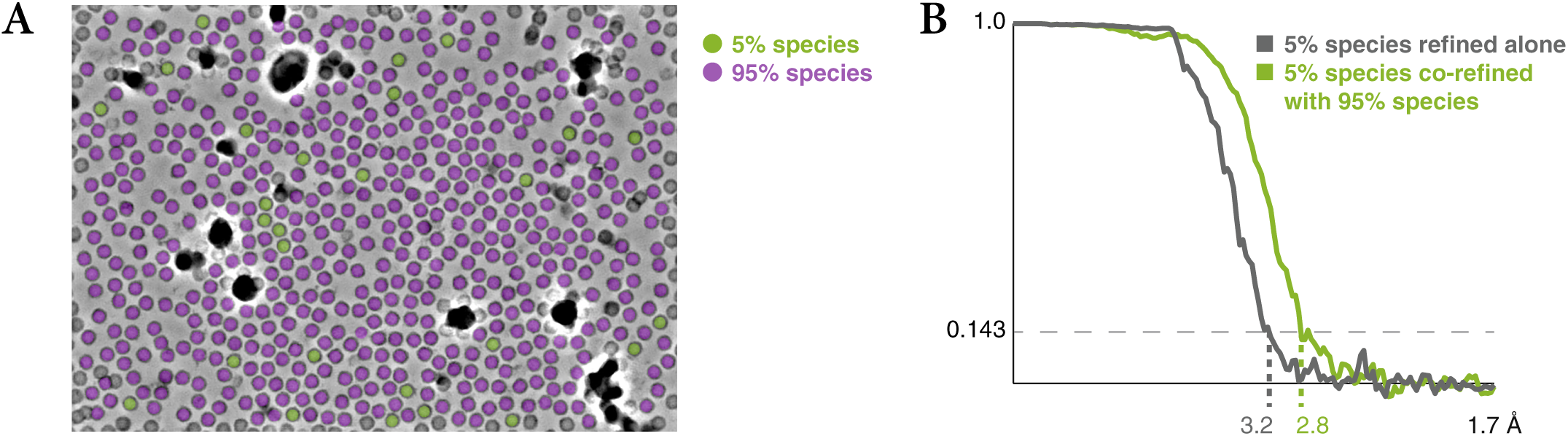
Benefits of considering more particles per micrograph through multi-species refinement. Apoferritin frame series were refined using a small 5% sub-population of the particles alone, and together with another 95% sub-population that improved the accuracy of the multi-particle system hyperparameters, but did not contribute particles to the 5% half-maps. (a) Exemplary distribution of the 2 sub-populations within a frame series. (b) FSC curves between the half-maps of the 5% population in both scenarios, showing the benefit of multi-species refinement.

**Figure S2.**
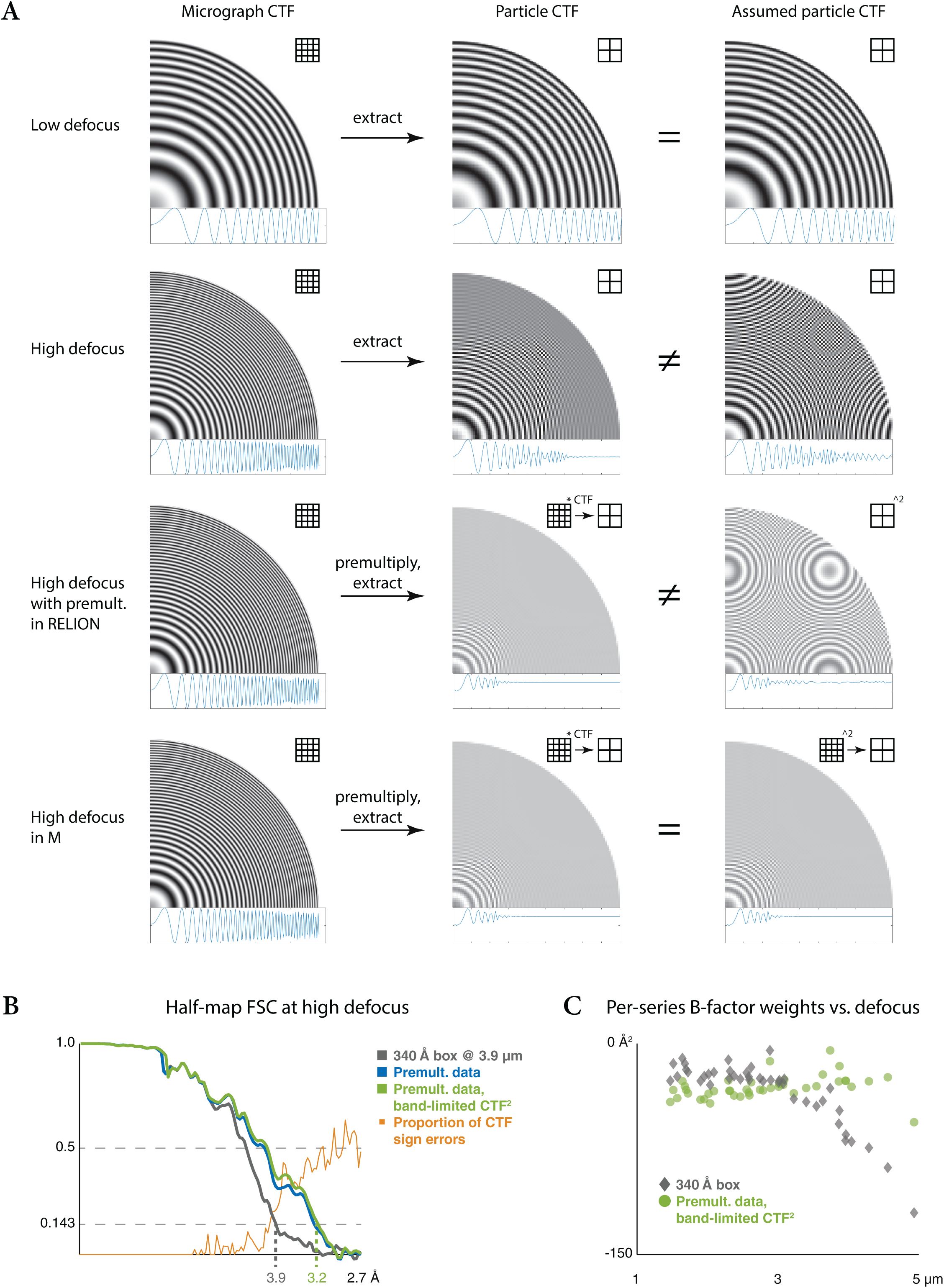
CTF correction in small particles boxes at low and high defocus. High-resolution information is delocalized at high defocus. Choosing an insufficiently large particle box size results in loss of that information. In Fourier space, this results in CTF oscillations becoming too fast to be resolved at the sampling rate provided by the small box, averaging to 0. M chooses the box size automatically for each frame or tilt series’ defocus, pre-multiplies the data and simulated CTF by the CTF to eliminate the oscillations and localize the signal, and then crops the data to the desired map size. This avoids the pitfall of losing map resolution due to an inappropriately chosen box size. (a) Visualization of the delocalization and aliasing effects in Fourier space as 2D and rotationally averaged 1D CTFs; grids depict sampling rate. At low defocus (row 1), all signal is localized within the box and no aliasing is seen in the simulated CTF used for the image formation model during refinement. At high defocus (row 2), high-resolution signal is delocalized outside the small particle box. Once the particle is extracted, the fast CTF oscillations are averaged to 0 and high-resolution information is lost. At the same time, the simulated CTF is filled with aliasing artifacts because it is not low-pass filtered in the same way. If the particle data are pre-multiplied by the CTF at a box size large enough to contain all signal and resolve all CTF oscillations (row 3), as can be done optionally in RELION, all particle signal is contained in the box after cropping it to a smaller size, and the CTF averages to 0.5. However, the simulated CTF^2^ does not match this and contains aliasing artifacts. M applies the pre-multiplication to both particle data and simulated CTF in a larger box before cropping (row 4) to avoid the mismatch. (b) FSC between the half-maps reconstructed from HIV1 virus-like particles of a single high-defocus (3.9 μm) tilt series in an insufficiently large box. Using data extracted without pre-multiplication, as is currently common, limits the resolution to 3.9 Å (grey). Pre-multiplying both particle data and CTF in a larger box, as automated in M, provides the best 3.2 Å result (green). Pre-multiplying only particle data is only slightly worse here (blue), but would likely lead to noticeably worse results in RELION as the aliased CTF^2^ would be used in the image formation during refinement. The FSC curves diverge as the proportion of CTF sign errors (orange) increases. (c) Relation between tilt series defocus and associated contribution of high-resolution information to the reconstruction. For the larger data set, not pre-multiplying the data results in a strong correlation, where high-defocus data is down-weighted to contribute less (grey). The correlation disappears when pre-multiplication is applied, so more tilt series contribute high-resolution information (green).

**Figure S3.**
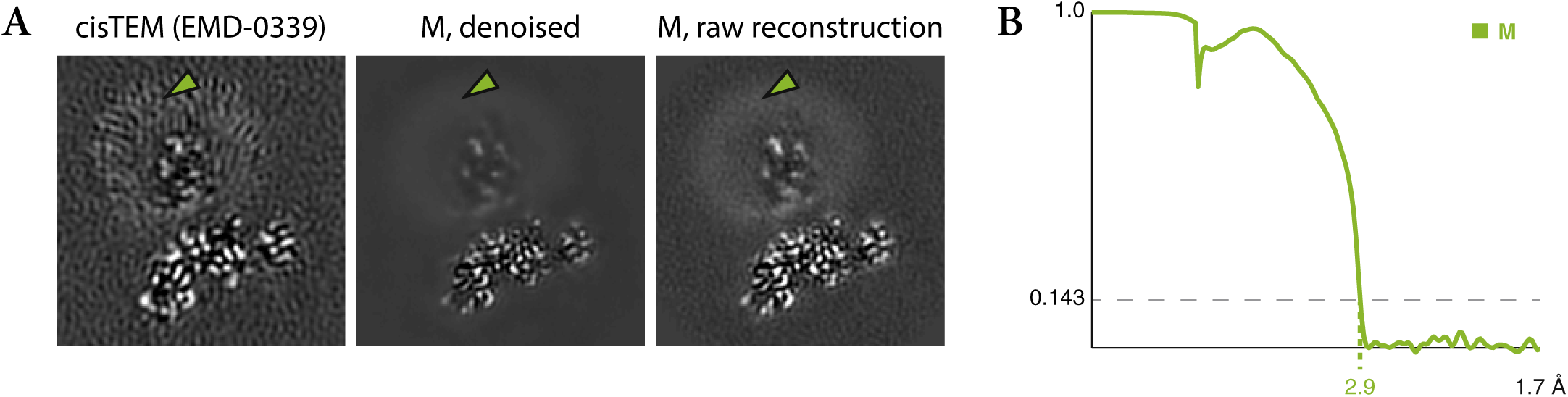
Effects of deep learning-based denoising of reconstructions during refinement. M trains a denoising model on each species’ half-maps after every refinement iteration to filter the maps to local resolution and avoid artifact over-fitting in low-resolution areas, such as lipid nanodiscs or flexible domains. (a) 2D XY slices through 3D reconstructions of the cannabinoid receptor 1-G membrane protein^28^. The original refinement in cisTEM (left) introduced artifacts in the highly disordered lipid region (green arrow). The denoised map (middle) and the raw reconstruction before denoising (right) used in the last refinement iteration in M are devoid of the artifacts because the denoising filtered and downweighed the low-resolution region. (b) FSC between the half-maps refined in M, showing a global resolution of 2.9 Å. A value of 3.0 Å was reported in the original study, with no FSC curve included with the deposited map.

**Figure S4.**
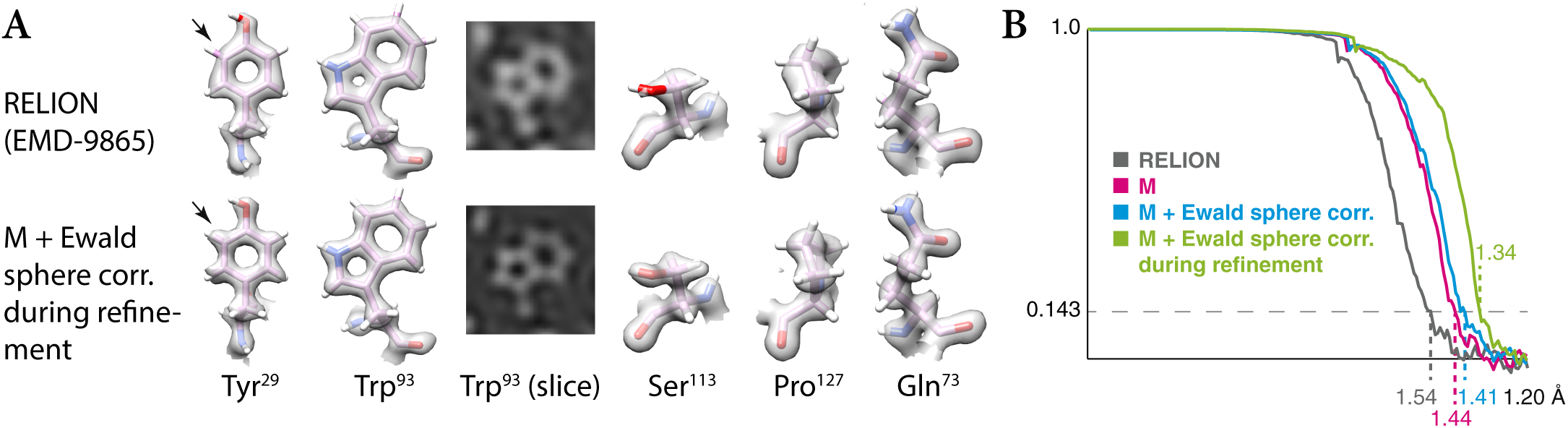
Comparison with RELION on atomic-resolution frame series data. Atomic-resolution data of apoferritin previously refined with RELION 3.1 to 1.54 Å (EMD-9865) were processed with M to achieve a resolution of 1.34 Å, showing that M’s image artifact model is suited for very high resolution. (a) Examples of side-chain densities produced by RELION (top) and M (bottom), showing cases of improved atomic features such as one of the hydrogens in Tyr29 (black arrow). (b) FSC between the half-maps produced by RELION (grey) and M (green), showing a general improvement in resolution through M.

**Figure S5.**
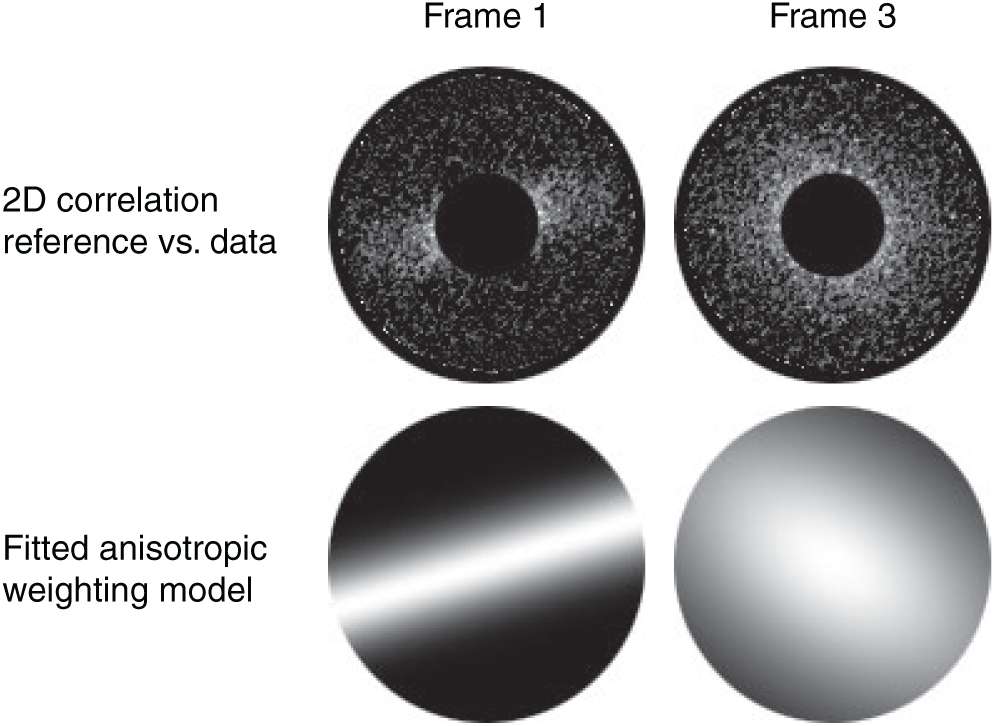
Examples of anisotropic B-factor weighting. Normalized 2D correlation between reference projections and data, averaged over all particles in a single frame is shown for the 1st and 3rd frame of the same exposure. Values in the low-frequency region are excluded to reduce the value range. The fitted B-factor is highly anisotropic for the 1st frame because of intra-frame motion: 0 Å^2^ and -62 Å^2^ along X and Y, respectively. For the 3rd frame, the fit is much more isotropic due to lack of intra-frame motion, but some high-resolution information is lost to radiation damage: -8 Å^2^ and -10 Å^2^ along X and Y, respectively.

